## supplementary Table 1 for "Adaptation of cyanobacteria to the endolithic light spectrum in hyper-arid deserts"

**Table S1**. Metadata and metagenome accession numbers for endolithic isolates

| Strain name | Origin | Substrate | Taxonomy | IMG ID |
|---|---|---|---|---|
| S-NGV-2P1 | Timna Park, Negev Desert, Israel | sandstone | *Chroococcidiopsis* | 3300039401 |
| G-MTQ-3P2 | Monturaqui, Atacama Desert, Chile | gypsum | *Chroococcidiopsis* | 3300037877 |
| C-VL-3P3 | Valle de la Luna, Atacama Desert, Chile | calcite | *Chroococcidiopsis* | 3300039404 |

**Table S2**. Gene distribution in the FaRLiP cluster of *Chroococcidiopsis* G-MTQ-3P2

| genome locus | function | gene symbol | annotation | protein size (aa) | gene size (bp) |
|---|---|---|---|---|---|
| Ga0395813_0004_87482_87718 | photosystem II PsbH protein | psbH2 | KO:K02709 | 78 | 234 |
| Ga0395813_0004_87766_89256 | photosystem II CP47 chlorophyll apoprotein | psbB2 | KO:K02704 | 496 | 1488 |
| Ga0395813_0004_89765_91165 | photosystem II CP43 chlorophyll apoprotein | psbC2 | KO:K02705 | 466 | 1398 |
| Ga0395813_0004_91234_92292 | photosystem II P680 reaction center D2 protein | psbD3 | KO:K02706 | 352 | 1056 |
| Ga0395813_0004_92384_92905 | FRL-allophycocyanin | apcD3 | KO:K02095 | 173 | 519 |
| Ga0395813_0004_92871_95210 | phycobilisome core-membrane linker protein | apcE2 | KO:K02096 | 779 | 2337 |
| Ga0395813_0004_95279_95758 | FRL-allophycocyanin | apcD2 | KO:K02095 | 159 | 477 |
| Ga0395813_0004_95873_96358 | allophycocyanin beta subunit | apcB2 | KO:K02093 | 161 | 483 |
| Ga0395813_0004_96460_96936 | FRL-allophycocyanin | apcA2 | KO:K02095 | 158 | 474 |
| Ga0395813_0004_97067_98065 | photosystem II P680 reaction center D1 protein | psbA3 | KO:K02703 | 332 | 996 |
| Ga0395813_0004_98412_99542 | photosystem II P680 reaction center D1 protein | ChlF | KO:K02703 | 376 | 1128 |
| Ga0395813_0004_99771_100004 | hypothetical protein | hypothetical | | | |
| Ga0395813_0004_100391_102259 | DNA-binding response OmpR family regulator | rfpB | COG0745 | 622 | 1866 |
| Ga0395813_0004_102532_105279 | light-regulated signal transduction histidine kinase | rfpA | COG4251 | 915 | 2745 |
| Ga0395813_0004_105417_105794 | CheY-like chemotaxis protein | rfpC | COG0784 | 125 | 375 |
| Ga0395813_0004_106087_106524 | CheY-like chemotaxis protein | cheY-like | COG0784 | 145 | 435 |
| Ga0395813_0004_106961_109309 | photosystem I P700 chlorophyll a apoprotein A1 | psaA2 | KO:K02689 | 782 | 2346 |
| Ga0395813_0004_109455_111677 | photosystem I P700 chlorophyll a apoprotein A2 | psaB2 | KO:K02690 | 740 | 2220 |
| Ga0395813_0004_112052_112603 | photosystem I subunit 11 | psaL2 | KO:K02699 | 183 | 549 |
| Ga0395813_0004_112614_112838 | photosystem I subunit 8 | psaI2 | KO:K02696 | 74 | 222 |

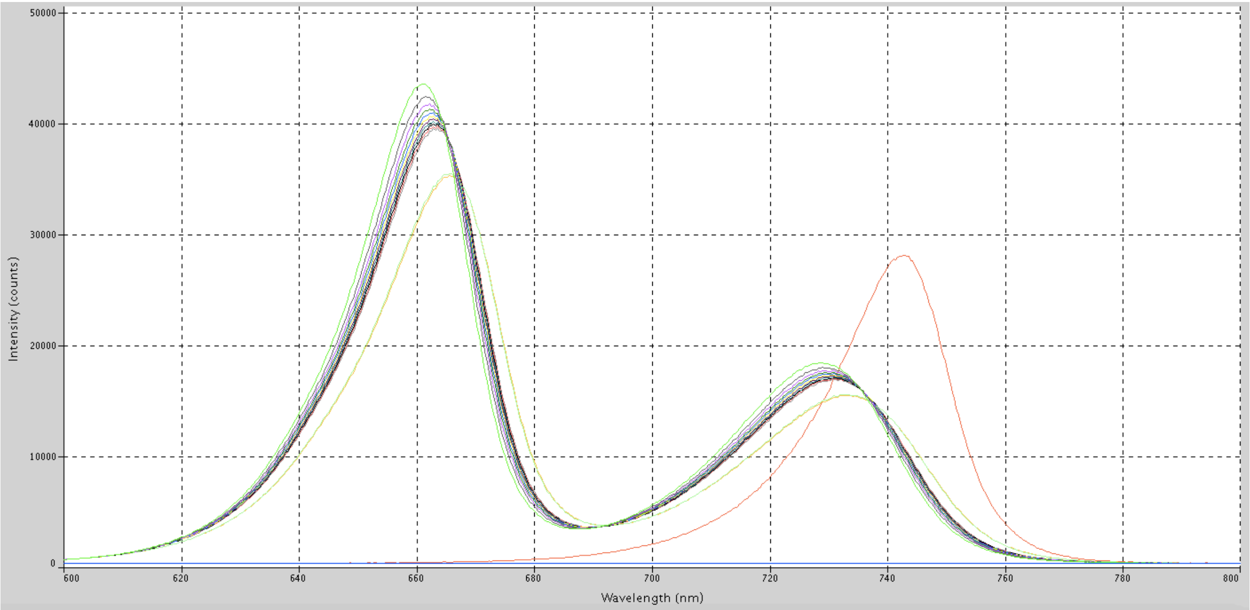


**Figure S1**. Spectra of white and far-red lights used in growth experiments.

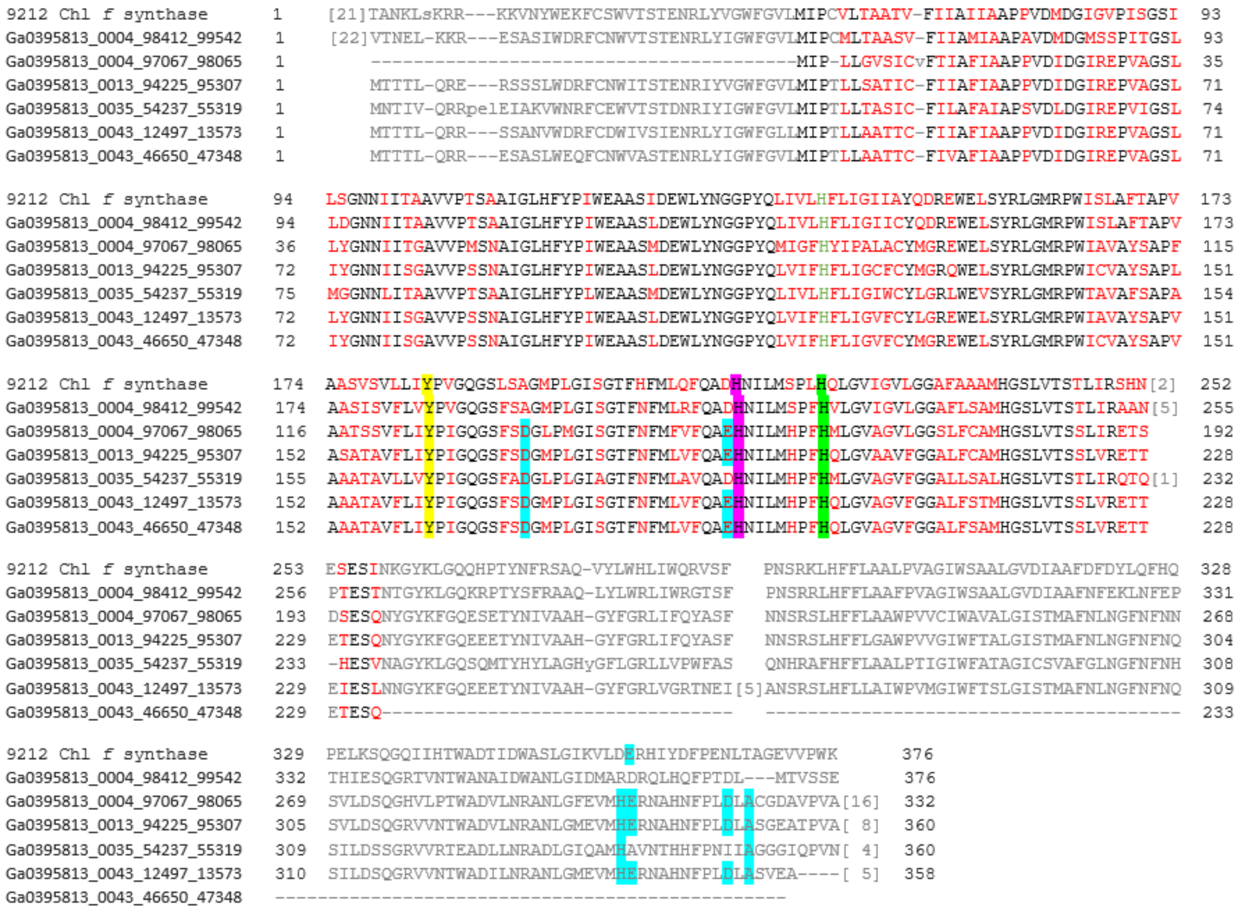


**Figure S2.** Protein alignment of Chl *f* synthase from *Chlorogloeopsis fritschii* PCC 9212 (9212 Chl *f* synthase) with Chl *f* synthase identified in the G-MTQ-3P2 (Ga0395813_0004_98412_99542), as well as all annotated *psbA* genes. Ligands to the $Mn_4Ca_1O_5$ cluster are highlighted in light blue. Tyrosine Yz residues are highlighted in yellow. Histidine residues thought to be involved in binding P680 Chl *a* are highlighted in green. Residues thought to ligate an additional Chl *a* are in green font. Residues involved in proton-coupled electron transport are highlighted in pink. Black residues are conserved across all sequences and red residues are not. Ga0395813_0004_98412_99542 lacks any ligands to the $Mn_4Ca_1O_5$ cluster, characteristic of Chl *f* synthase [17].
